# A modeling algorithm for exploring the architecture and construction of bird nests

**DOI:** 10.1101/600718

**Authors:** Hadass R. Jessel, Lior Aharoni, Sol Efroni, Ido Bachelet

## Abstract

Natural biological structures are often complex and cannot be mapped directly to genes, being therefore impossible to explore by traditional biological tools. In contrast, digitizing these structures enables to explore their properties and behavior under specific conditions, by means of computational manipulations, simulations, and analyses. We describe a generic algorithm for the digitization and exploration of the complex structures exhibited by common, interwoven bird nests. This algorithm takes as input computerized tomographic scans of the studied Dead-Sea Sparrow (*Passer moabiticus*) nest, identifies and isolates each branch entity within the three-dimensional data and finally extracts the characteristics of each branch. The result is a reliable three-dimensional numerical model of the nest that contains a complete geometric dataset per each of its components, e.g. dimensions and contact points with neighboring components, as well as global properties, e.g. density distribution and network structure. Based on these, we were able to simulate various models of the nest construction process. Altogether, the described algorithm and possible derivatives thereof could be a valuable tool in studying the structure-function relationships of similarly complex biological objects.

## Introduction

The field of structural biology is concerned with the relationships between structure and function of biological objects, mainly at the molecular level. Testing specific hypotheses regarding these relationships is typically done by altering the structures using traditional experimental techniques such as genetic mutations, and measuring the consequent behavior of the object under study^1^. However, complex biological structures at the macro-scale level, such as animal made structures, e.g termite mounds^2^, orb webs^3^, and bird nests^4^, that presumably have a genetic basis, are challenging and often impossible to explore this way, since there is no simple injective mapping between genotype and phenotype.

While such macro-scale biological structures cannot be studied by traditional biological tools, their structure-function relationships can be studied *in-silico* in digital formats. Digitizing complex structures by converting the objects using three dimensional (3D) imaging techniques, such as 3D scanning, computer tomography (CT), and magnetic resonance imaging (MRI) into numerical (digital) model representations, enables to explore their properties and behaviour under specific conditions, by means of computational manipulations, simulations, and analyses^5–8^. These structures may be 3D printed prior to generating digital manipulations to further explore the structure, e.g by means of mechanical testing^9^. To obtain numerical representation in form of 3D solid or surface models from medical imaging modalities, a computer-based image reconstruction is carried out on a set of two dimensional (2D) tomograms. Digital analysis of the data and 3D models obtained from the actual biological structures provides us with a definitive description of the studied structures.

Computational modeling has greatly augmented our understanding of the role of structure on biological function, and is now a growing strategy for biomechanics and medical research, with recent advances in model development and simulation platforms leading to growing contributions in medicine and comparative biomechanical studies^10^. The popularity of digital analyses may be due to many factors, from their ability to provide robust quantitative analyses for complex and intricate structures often found in biological systems, to the ability of numerical methods such as Computational fluid dynamics (CFD) and Finite element analysis (FEA) to provide colossal amounts of output data, allowing robust statistical analyses and detailed graphical displays.

Digitized structures enables us to simulate and predict how a structure will perform in natural or under changed operating conditions and investigate the effect of induced structural modifications on the natural function. For example, frontal sinuses in goats and other mammals are assumed to function as shock absorbers, protecting the brain from blows during intraspecific combat. Generating 3D models of domesticated goat (*Capra hircus*) skull with variable frontal bone and frontal sinus morphologies and simulating loading scenarios using the finite element method (FEM) shows how morphological modifications affect distribution of stresses in the skull^11^. Another study shows the effects of deteriorated trabecular bone structure on bone stiffness and strength by generating healthy 3D printed trabecular bone model and comparing it with the same model after bone resorption was simulated. Since the deteriorated structural bone model is derived from the healthy one, it is possible to directly estimate the decrease of tissue stiffness and strength as a result of bone resorption for the specific structure^12^. These studies show how digitizing biological structures is beneficial for understanding and evaluating the performance of complex natural structures.

The physical structure of biological structures often reflects their assembly and function. Bird nests are no exception, containing numerous interwoven components with structural stability of the architecture depending mainly upon the structural organization of the components and their material properties. Avian nests are essential for reproductive success, providing a location for rearing the young and safely supporting the incubating bird and its clutch of eggs^13^. Nests have been shown to be constructed out of a wide variety of organic and artificial building materials^14^, which can generally be classified as being either structural materials or lining materials. While structural materials make up the general shape of the nest and provide structural support for the parents and offspring, lining materials generally create a suitable microclimate for raising offspring^15,16^. It is assumed that building materials within different parts of a nest are not randomly selected and appear to have a structural role^17^. Recent studies have tried to determine the construction patterns of nests and the factors that affect nest construction both examining the building materials and their mechanical properties^17^. Studying nest construction is primarily performed by means of deconstruction and characterization of its components to relate the composition of the nest regions to their function^17,18^. Consequently, someone who is interested in rebuilding the structure or performing mechanical tests to study its biomechanics properties, will likely be at a disadvantage. Developing *in-silico* nest models based on actual nest structures can be labor intensive and time consuming, nevertheless the change from physical to digital data has made possible to perform numerous experiments and model modifications. Digitizing scanned nests and classifying each branch entity, can be done with semi-automated and manual segmentation approaches. Yet, this approach is not feasible due to similarity in material properties of branch components, which results in a challenging and time-consuming task of separating between nest components.

The goal of this study is to design and implement an algorithm that is capable of precisely mapping the structure of interwoven birds nests, a relatively complex task due to the intricate multipart structure. In this method we take CT scans of interwoven birds nests as an input, identify the nest structure within the 3D data using image processing and thinning techniques. The novel algorithm is then used to isolate each branch entity. Finally the characteristics of each branch are extracted and used for statistical analysis of the structure and for generating graphical visualizations and simulations of construction patterns. Better understanding the relationship between structure and function can be done by breaking down the nest into the elements comprising it and studying how each branch affects the architecture as a whole. *Mutating* the nest structure by elongating, trimming, or removing components, results in new models that can be used in comparative simulations for studying the structure stability and investigate bird nests building techniques.

## Results

We used Dead-Sea Sparrow (*Passer moabiticus*) compound nests built by assembling branch components (**Fig. 1A**). The studied Dead-Sea Sparrow nests are relatively large, assembled by thick elements with a mean branch thickness of 2.6mm, being an advantage in the following computational analysis. A case study was carried out to illustrate and evaluate the proposed method. **Figure 1B** shows a schematic overview of the method. This algorithm was designed in a pipeline pattern, starting from the image sequence, passing through different filters to achieve the final nest numerical model. In each step and sub-step (**Supplementary Fig S1**), the intermediate result is written to a file, to allow easy management of the complex and computationally-intensive process.

**Figure 1.**
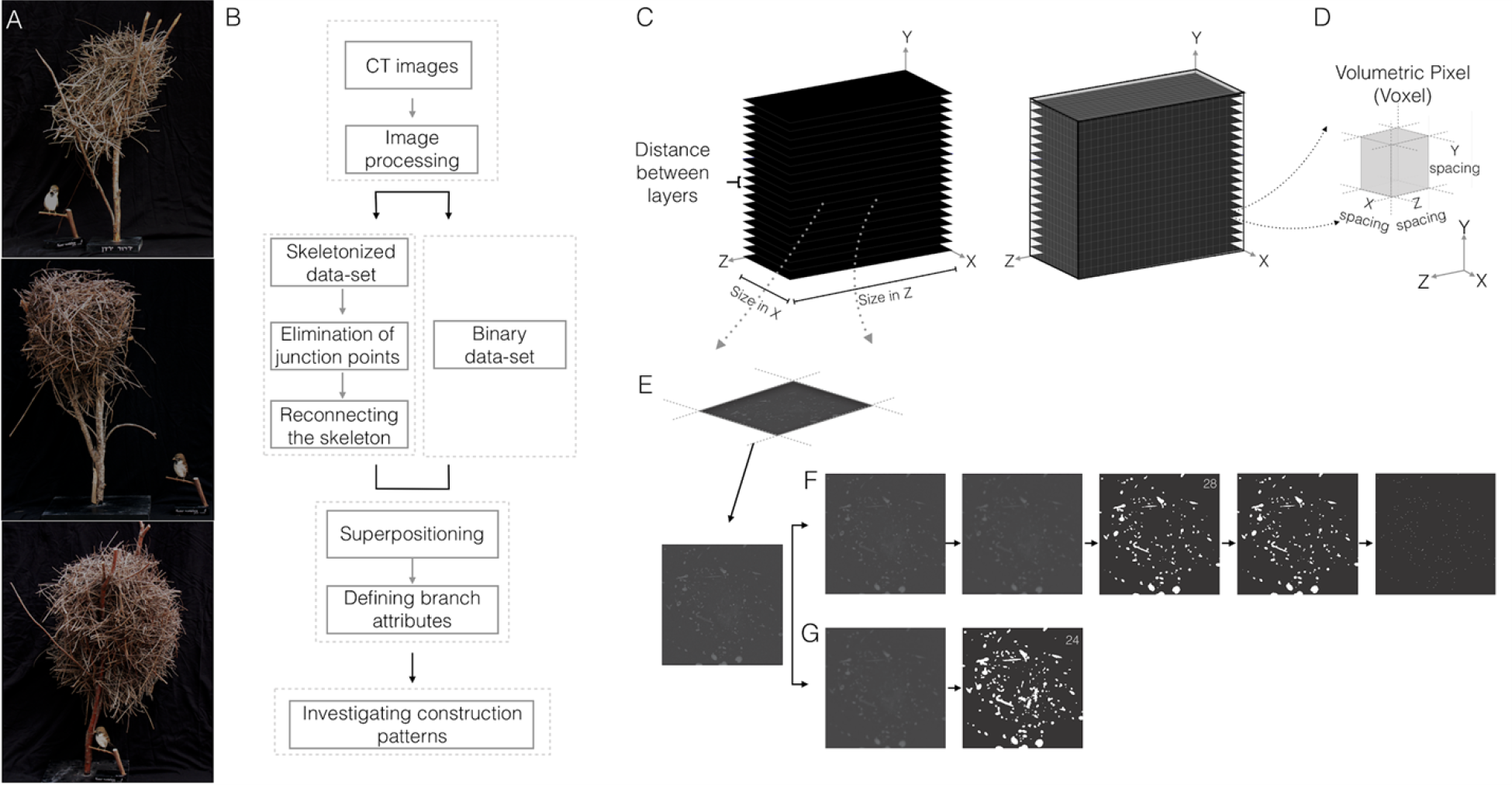
CT image data processing workflow and representative nests. (A) Dead-Sea Sparrow (*Passer moabiticus*) representative nests. (B) Schematic overview of the modeling algorithm. The algorithm first operates on the original CT scans of the studied nest, to standardize and normalize the input. The first stage yields two image types, raw skeleton and binary datasets. A dedicated algorithm is applied to the skeleton 3D model eliminating all junction points between edges, then reconnecting highly correlated edges that originate from the same branch. Superpositioning the binary data set and skeleton is used to calculate branch thickness, length and contact points. Construction patterns of nests can then be investigated (C) CT image dataset is generated, showing a voxel intersecting a layer (D). (E) Layers are processed in parallel in Fiji, (F) Showing image processing of grayscale layers; grayscale after neighbourhood averaging 3D filter in the z axis (□=5), 3D gaussian blur filter (□=1), and global thresholding with a value cut off of 28. Next, a hole filling operation is applied and dataset is skeletonized to identify the three dimensional centreline of the nest structure. (G) A second dataset is processed using a gaussian blur filter (□=2) and a global thresholding cut off value of 24.

To digitize and explore the complex structures exhibited by common interwoven bird nests, we used our method to optimally take computerized tomography (CT) scans of the studied nests as an input, and identify and isolate the 3D structure. This method first analyzes the greyscale CT derived datasets using a slice-by-slice segmentation process to identify potential branch locations in the 2D cross-sectional images (**Fig. 1C**). Complete structural 3D information of the nests is obtained by X-ray computer tomography set to an in-plane resolution of 512 × 512 with approximately 3700 scan slices available with non isometric voxels (i.e., the voxels are cuboids instead of cubes) (**Fig. 1D**). The resolution was found to be sufficient for analyzing the individual nest components and identifying the nests’ architecture. CT datasets are imported into FIJI^19^ with a background type of 8-bit unsigned integer (**Fig. 1E**). The datasets are preprocessed using image processing filters, which are used for noise reduction and averaging. The filters include a neighbourhood averaging 3D filter in the z axis (□=5), and a 3D Gaussian Blur filter (□=1) (**Fig. 1F**). Following preprocessing a new datasets was generated for yielding the structure skeleton. Since branches appear in a considerable variation of diameters (1.2 mm to 8.8 mm) a range of intensity thresholds are tested to optimally identify all branches. The optimal threshold cut off value is one that reduces image noise while preserving maximum significant image data (T=28) (**Supplementary Fig. S2**). As this image sequence is next skeletonized, it is crucial to eliminate inner branch cavities to ensure that the skeletonization process would not fail to describe the actual branch geometry. Thus, a hole filling operation is applied for filling branches with inner voids or small hollow regions due to local image noise. Finally, a 3D thinning algorithm is applied^20,21^ to obtain the structure’s skeleton, from which the 3D centerlines of individual branches are to be identified.

The thinning process does not identify touching branches as separate entities. Rather, in the resulting skeleton, branch centerlines may be connected with bridges that originate from contact regions between branches (**Fig 2A**). **Figures 2A-D** show the approach as applied to a sample of branches. A generalized method was designed to identify *true* and *false* branches. The algorithm was developed to optimally break the entire skeleton structure into simple edges by eliminating all junctions and forks. The skeleton is first cleaned by pruning tail edges, using an iterative elimination process with a predefined length threshold (T=5mm). Next, all junctions are identified (black arrowheads) and pruned out of the structure, this is done by removing pixels (equivalent to 2mm) off each junction edge, resulting in a simplified structure that contains simple edges only (no forks). As this process eliminates both *true* and *false* junctions, it is necessary to reconstruct the *true* junctions. Various separation and connection scenarios were considered for the development of the algorithm (**Supplementary Fig. S3**). A matching algorithm is then applied to rate the correlation between endpoints *p* of branches oriented in a predefined proximity (**Fig. 2E**). To systematically assess whether two segments originate from the same branch, we calculate branch friendship score *C* via

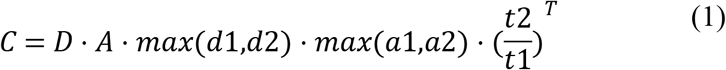

**Figure 2.**
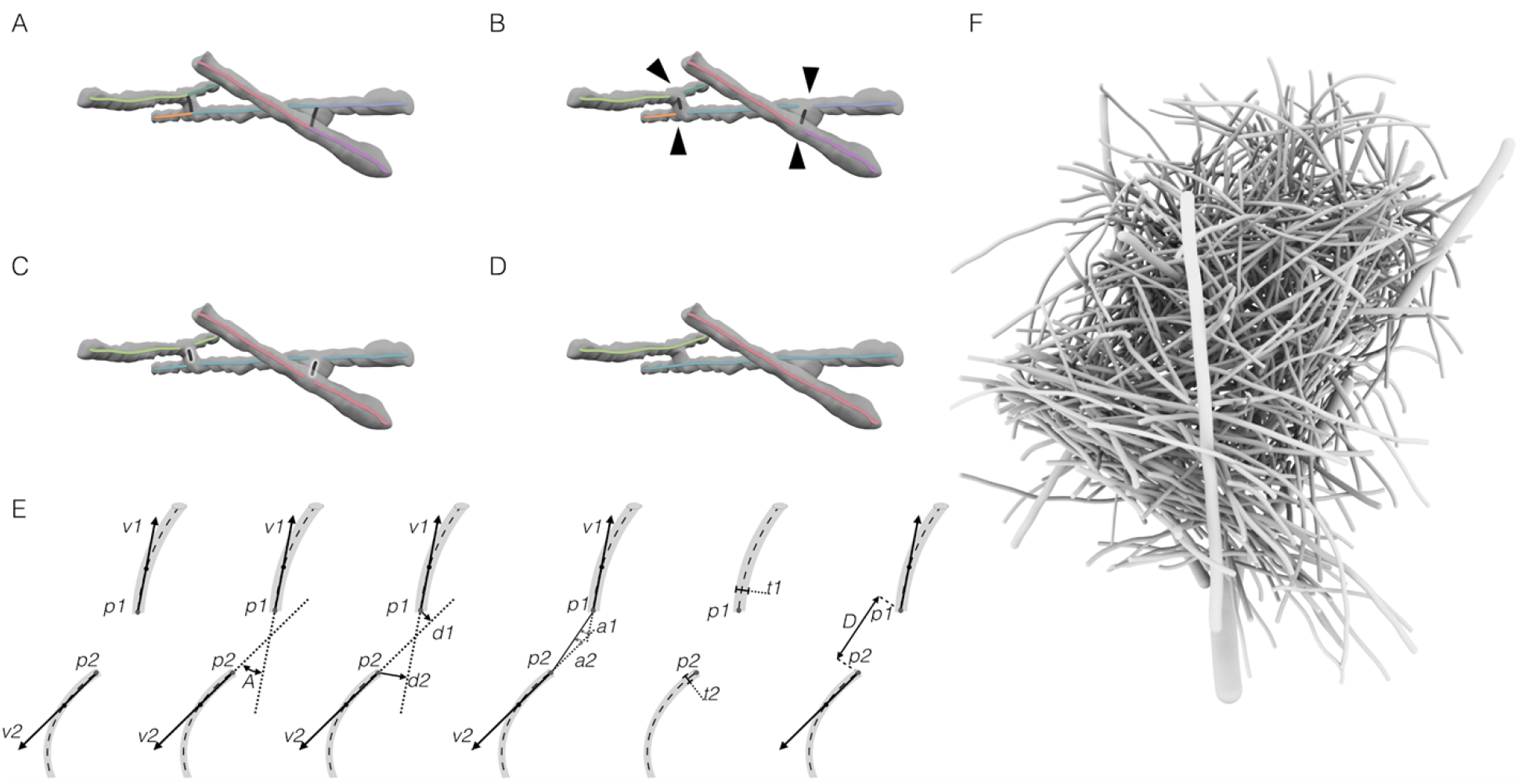
Digitizing an avian nest structure. Illustration example of the process of breaking the skeleton structure and reconnecting highly correlated edges. (A) Structure skeleton with branches represented as edges with different colors (n=9), including 4 junction vertices and 6 end vertice. (B) Breaking the structure by eliminating junction points (black arrowheads), the resulting structure is composed of end-end edges (n=9). (C) Edge groups with high inter-correlation are marked with the same color, with correlated edges linked to each other through the high correlation endpoint couples. (D) Short edges representing the contact points are eliminated. (E) Illustration showing the different parameters used to measure the correlation between end-points of two branches. Two end-points noted by *p1* and *p2*, The direction of the branch at each end point is measured as a vector from the end point to a point along the branch in a constant length T measure along the branches skeleton. The two direction vectors are noted by *v1* and *v2* respectively. The angle between the two direction vectors is noted by *A*. *A* measurement of the distance between each direction vector and the end point on the other branch is used, the two distances are noted by *d1* and *d2* corresponding to *p1* and *p2*. Measurement of the angle between each direction vector and the vector formed by the two points, noted by *a1* and *a2*. Thickness of each branch at the corresponding end point is noted by *t1* and *t2* respectively. The thickness is measured as the median taken over a fixed number of sample points on the branch’s edge. The distance between *p1* and *p2* is noted by *D*. (F) 3D model of reconstructed nest.

Where *D* is the distance between the endpoints, orientation difference *A* is measured as the spatial angle between the orientation vectors *v* withl80 – 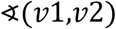, in which an angle of zero represents a perfect orientation match. The distances *d1* and *d2* are measured between *p1* and *v1*, and *p2* and *v2* respectively. Angles *a1* and *a2* are measured between the vector *p1→p2* and *v1* and *v2* respectively. Finally, the thickness *t* of each branch is recorded. The friendship score is therefore inversely related with correlation between two endpoints. These parameters often exhibit non-proportional effect on the score, requiring an enforcement of cutoff values before plugging them into the formula. If a parameters value is smaller than a certain minimum it is replaced with the predefined value. For *D* and *d* minimum values of 1.0 and 0.5 were enforced respectively. The value enforced on *t2/t1* is 0.1 and *T* is 3. The actual implementation of the scoring formula can be seen in **Supplementary Fig. S4**. Highly correlated branches are iteratively linked combining two edges into a single edge (**Fig, 2C**), and finally short edges are removed (**Fig. 2D**). This is based on the assumption that highly correlated edges originate from the same branch. The preliminary result of the python code is a list of skeletonized branches, where each branch is described as a set of voxels.

To calculate branch thickness we used an additional binary data set derived from the original CT scan. The original dataset is preprocessed using a 2D Gaussian Blur filter (□=2) (**Fig. 1G**). Testing a range of threshold intensities reveals an optimal cutoff threshold value (T=24), that provides binary images from which branch thicknesses can be extracted. The threshold cutoff value is selected by measuring the thickness of several branches and comparing the diameters to scanned images after thresholding, therefore obtaining true thickness values. Branch thickness is analyzed at each skeleton point, by superpositioning the skeleton and the binary image sets. Measured diameter is then added to each voxel unit. An algorithm is applied to clean out branches with zero thickness, and assign thickness to short branch segments with zero thickness. Finally, a constant thickness value is documented for each branch by calculating the median thickness of sample points along the branch.

Contact points between branches are identified by measuring the distance between branch surfaces, as the distance between the skeleton points and branch diameter is known. We were able to efficiently produce visualization of the reconstructed nests and qualitatively assess the accuracy of the reconstruction done by the algorithm as well as print 3D models of the nests **Supplementary Fig. S5**. This analysis reveals both visually and quantitatively the distribution of twig length, thickness and degree of connectivity across different parts of the nest (**Fig. 3A-D**). The findings show that the nest is comprised of 739 branches with a large contact network of 208 intersection points, connecting the branches in the structure (**Fig. 4A**).

**Figure 3.**
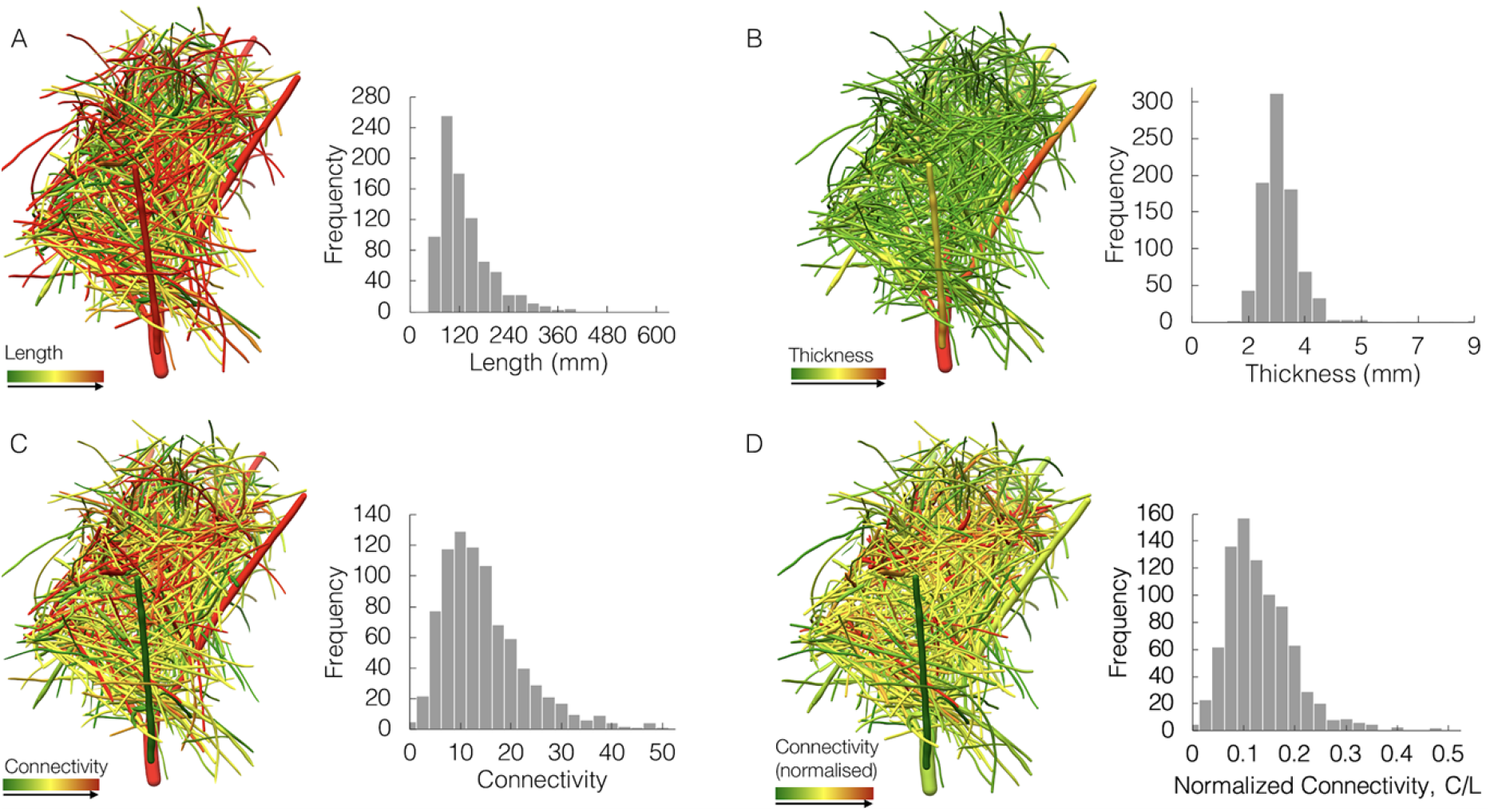
(A) Branch length, with low length in green and high in red (left). Frequency of branches for different length values (right). (B) Branch thickness, with low thickness in green and high in red (left). Frequency of branches for different mean thickness values (right). (C) Branch connectivity, with low connectivity in green and high in red, showing connectivity of each branch in a uniform color (left). Frequency of branches for different connectivity values (right). (D) Branch normalized connectivity, with low connectivity in green and high in red, showing normalized connectivity of each branch in a uniform color (left). Frequency of branches for different normalized connectivity values (right).

**Figure 4.**
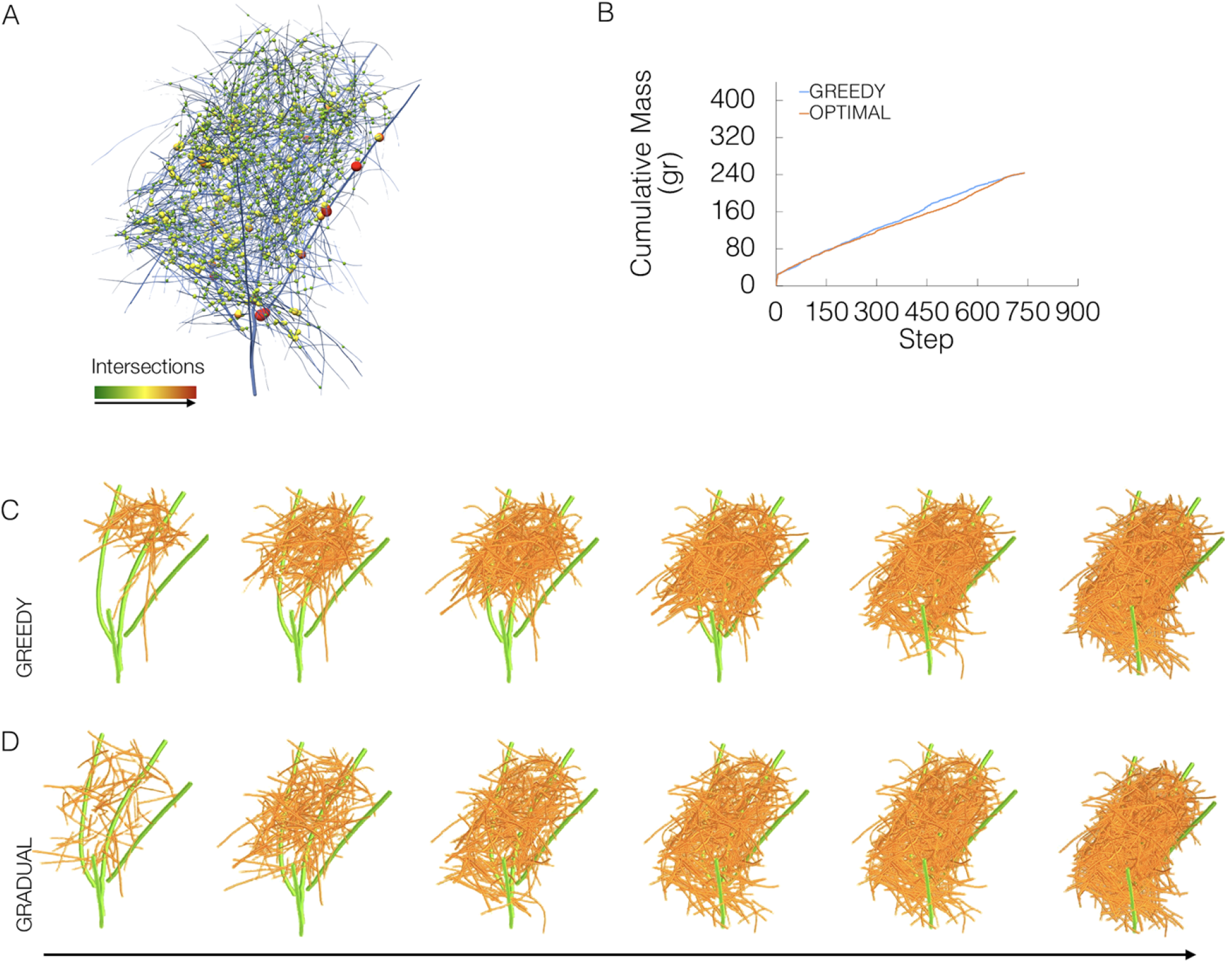
Nest intersection points and construction patterns. (A) Skeleton proportional to original thickness (25% of original thickness) showing intersection points as spheres. Color and radius of spheres is proportional to the thickness of branches intersecting. (B) Graph represents the cumulative nest mass per step, comparing the two construction strategies. Where in each step an estimated mass of the newly inserted twig is added. (C-D) Series of three-dimensional nest models showing two building algorithms to test alternatives for building avian structures. (C) GREEDY algorithm first identifies the structures scaffold (green), then each branch in turn is added, when in each step the branch that is most gravity-connected is added to the existing structure. (D) GRADUAL algorithm first builds a minimum viable skeleton that can hold the structure, that is further enriched with branches.

To investigate variations in construction patterns, we used different building algorithms to test alternatives for constructing avian structures. Our method describes the nest as a directed network, where centers of branches are marked as spheres. Spheres size correlates with branches diameter and edges connecting between spheres resemble contact points between branches (**Fig. 4A**).

Based on this result, we created a first algorithm, GREEDY, that automatically identifies the contact points from the contact network and assigns each branch with a value—*gravity-connected*—defined by the magnitude of the link between branches and their relationship to the gravity vector. GREEDY first identifies the structures scaffold, usually the thickest branches originally attached to the tree. Each branch in turn is added, when in each step the branch that is most *gravity-connected* (i.e. the most structurally supported branch by the already constructed structure) is added to the existing structure. A second algorithm was then created, GRADUAL, that automatically identifies the contact points from the contact network and assigns each branch with two values, *gravity-connected* and *gravity connectivity*. GRADUAL first identifies the structures scaffold. Each branch in turn is added, when in each step the branch that is least *gravity-connected* and has *gravity-connectivity* to stay put due to friction with other branches, is added to the existing structure. GREEDY and GRADUAL reach the final nest in very similar trajectories in terms of cumulative nest mass (**Fig. 4B**). However, in GREEDY, the branches in the center of the nest, that have several contact points, are first to be placed, followed by the external branches (**Fig. 4C**). In contrast, GRADUAL results in a more complex process that first builds a minimum viable skeleton that can hold the structure, that is further enriched with branches (**Fig. 4D**).

## Discussion

Despite the advancement in computational modeling and numerical analyses tools available to date, simulation studies overall adoption rate have been limited, in large part, by the challenges associated with the translation of medical images into numerical models. The workflow described here solves problems that constrain traditional computational modeling approaches, mainly regarding time costs with segmentation and computationally limited file size. Image semi-automated and manual segmentation approaches can be time consuming especially when studying complex structures such as interwoven bird nests which have multiple parts that require a subjective segmentation assessment of where each region of interest starts and ends. Moreover, this algorithm, which is designed in a pipeline pattern where in each step and sub-step the intermediate result is written to a textural file and read back before the following step, allows an easy management of the complex and computationally intensive process. The translation of image files into a textual format allows access to the data in a reasonable amount of time, and lowers the complexity of both in/out operations and memory consumption.

Here, we describe a dedicated generic algorithm that successfully dissects and reconstructs a multipart object, and directly translates it into a numerical model. The method described and implemented herein points towards new modeling opportunities for which the barriers between the physical and digital domains can be eliminated, enabling the digital visualization, analysis and manipulation of complex, macro-scale biological structures, such as bird nests. The resulting models closely resemble the physical analog, making this process valuable for data analysis and computational biomechanics research. It is thus likely that scientific modeling tools in the future will incorporate a method similar to the one described herein, enabling scientists to access, digitally modify and analyze complex natural structures. Furthermore, in the future, advanced capabilities to convert CT scans of biological structures into digital data such as the one demonstrated herein may allow for sharing and exploration of diverse biological structures in the scientific community^10^.

## Methods

The proposed method consists of the following main steps: *branch segmentation, skeletonization*, and *branchpoint pruning and matching*. Each of these steps is described separately in the following subsections. In general the framework developed is based on first generating a digital model of the nest, and the separating between its component. The analysis results in a file listing the branch coordinates, their length, median thickness and contact points.

### Nests

Dead-Sea Sparrow (*Passer moabiticus*) compound nests (n=3) were obtained from the Steinhardt Museum of Natural History at Tel-Aviv University.

### Data

Nests were fixed for scanning in a dual-source CT scanner (Somatom Definition. Flash; Siemens Healthcare, Forchheim, Germany) system. The nests were scanned with the X-ray tube voltage set to 120 kVp and the X-ray tube current of 33 mAs. The best contrast was achieved using a sharp convolution kernel filter (V90μ) during the scans. X-ray projections were set to an in-plane resolution of 0.5mm and slice-to-slice separation of 0.1mm.

### Branch segmentation

The branch segmentation takes advantage of the relatively high contrast in CT images between the nest branches and surrounding air. A 3D Gaussian Blur filter is applied on each image slice to reduce image noise. Two new datasets are generated by setting different thresholding parameters, this is realised by setting an upper limit of allowed difference in voxel grey value. Next, a hole filling operation is applied for filling branches with inner tunnels and refining inner branch regions that may have small holes due to noise. It is crucial to eliminate inner branch cavities to ensure that the coming thinning process would not fail to describe the actual geometric structure. After branch segmentation two new binary sub-volumes are formed that represent the extracted nest; one that will yield the structure skeleton and another for the defining of branches volumetric data.

### Skeletonization

A binary nest formed in the previous step is skeletonized to identify the three dimensional centreline of the structure and to determine the branchpoint locations. Branches in the nest are interwoven and exhibit similar image characteristics which makes the nest appear as a single part. Hence, this process is key for defining and separation each branch entity. To obtain the skeleton, a sequential 3D thinning algorithm implemented in Fiji is used. A thinning function deletes border voxels that can be removed without changing the topology of the nest. The thinning is performed symmetrically and the resulting skeleton is guaranteed to lie in the middle of the cylindrically shaped branch segments.

### Branchpoint pruning and matching

After completion of the thinning step, the skeleton is smoothed by pruning false branches, and locations of the branchpoints are identified and pruned, resulting in simple edges without forks. The goal of branchpoint matching is to find topological correspondence between edges that originate from the same branch. A generalisation of different separation scenarios lies on the relative orientation of the branches and correlation between them. Therefore, various separation and connection scenarios were considered for the development of the algorithm, where edges are considered correlated based on the orientation, distances, and branch thickness of endpoints. Correlated edges are connected and branch parameters are calculated.

### 3D Printing

Model was printed on a Stratasys 0bjet500 Connex3 3D printer using a VeroWhitePlus material (RGD835).

## Acknowledgements

HRJ, LA, IB and SE wish to thank Daniel Berkowic 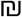 Assaf Uzan from the Department of Zoology, Tel Aviv University, The Steinhardt Museum of Natural History, for help with nests used in this study; Adva Sharabi, Ayelet Eran, & Aliza Turgeman from the CT Unit, Medical Imaging Department, Rambam Health Care Campus Haifa, for valuable technical assistance; and to Bnaya Bauer for valuable assistance and discussions.

## Author contribution

HRJ and LA designed the algorithm and the simulations; All authors analyzed data and wrote the manuscript. IB and SE oversaw research.

## Additional information

All the authors declare no conflict of interest.

